# Respiration shapes the neural dynamics of successful remembering in humans

**DOI:** 10.1101/2025.04.17.649286

**Authors:** Esteban Bullón Tarrasó, Fabian Schwimmbeck, Marit Petzka, Tobias Staudigl, Bernhard P. Staresina, Thomas Schreiner

## Abstract

Respiration has been shown to impact memory retrieval, yet the neural dynamics underlying this effect remain unclear. Here, we investigated how respiration shapes both behavioral and neural expressions of memory retrieval by re-analyzing an existing dataset where scalp electroencephalography and respiration recordings were acquired while participants (N = 18) performed an episodic memory task. Our results unveil that respiration influences retrieval-related power fluctuations in the α*/*β band and concomitant memory reactivation. Specifically, we found that both key neural signatures of successful remembering were co-modulated during exhalation, with the strength of the interaction between respiration and reactivation processes being associated with memory performance. Together, these findings suggest that respiration may act as a scaffold for episodic memory retrieval in humans by coordinating the neural conditions that support effective remembering.

**Significance statement:** Recent evidence suggests that respiration may shape neural dynamics underlying various cognitive processes. In this study, we identify respiration as a potential pacemaker for memory retrieval by showing that key neural signatures of effective remembering—namely, decreases in α/β power and the reactivation of previously encoded neural representations—are tightly synchronized with the respiratory cycle. Notably, the strength of this respiration-brain coupling is associated with individual memory performance, underscoring the critical role and functional significance of brain–body interactions in supporting cognitive functions.

## Introduction

Compelling evidence from both rodent and human studies demonstrates that breathing might serve a purpose beyond supplying oxygen; it shapes neural activity during wake^1,2^ and sleep^3,4^. Breathing-locked neural oscillations have been observed across diverse brain areas, oscillatory frequencies, and species^5–9^. While the precise mechanisms enabling this modulation remain unclear, evidence suggests that mechanoreceptors in olfactory sensory neurons, which respond to airflow, generate respiration-locked oscillations in the olfactory bulb^9–12^. These oscillations propagate along the olfactory pathway via the piriform and entorhinal cortices to neocortical and hippocampal regions^1,3^. However, since respiratory rhythm generators in the brainstem may also contribute to the entrainment of breathing-related neural oscillations^3,13–15^, the preferred pathway through which breathing modulates neural activity remains uncertain.

Nevertheless, respiration has been shown - potentially through its impact on neural activity - to influence perceptual^16^, motor^17,18^, and cognitive processes^19^ in humans. Performance is typically enhanced when stimuli are presented during inhalation^19–22^, suggesting that inhalation-linked neural processes may aid perception. In contrast, mental actions such as motor and visual imagery tend to occur during exhalation^18^, indicating that different phases of respiration may create time windows of opportunity for distinct cognitive functions^9^. Notably, memory processes have also been shown to be affected by the phase of respiration^23^. It has been demonstrated that retrieval-related reminder cues are more likely to trigger successful remembering if presented during inhalation as compared to exhalation^20^, potentially due to stimulus processing being facilitated during inhalation. However, it is unknown whether memory-related neural processes, aside from stimulus-driven effects, might be associated with respiration, particularly during later phases of the breathing cycle, such as exhalation. Altogether, the results reported above suggest that respiration, and specifically its phase, may directly influence our ability to remember. But how could respiration facilitate effective memory retrieval?

In humans, memory retrieval has long been associated with characteristic patterns of neural activity. For instance, low-frequency power decreases in the α/β range (∼8 – 30 Hz) have been consistently found to accompany successful remembering in cortical areas and the hippocampus^24–27^. The magnitude of those decreases has been proposed to reflect heightened levels of information processing and retrieval-related engagement of cortical modules^25^. In line with this assumption, it has been shown that α/β power fluctuations do not only track the quantity but also the level of detail of memory-related information represented in the brain^28–30^. Moreover, several studies have established a tight link between α/β power decreases and the emergence of retrieval-related memory reactivation^31,32^. Specifically, retrieved memory information seems to be particularly decodable from electroencephalographic (EEG) signals during low α/β power levels^28,33^, while other studies indicated that the low frequency oscillatory phase itself might carry a temporal signature of the reactivated memory content^34,35^. Importantly, beyond its close relationship with memory processes, α/β activity has been demonstrated to be robustly modulated by respiration during diverse cognitive tasks (i.e., visuospatial, lexical, and math)^19^, perception^16^ and rest^6^. Hence, respiration might accomplish its influence on the behavioral aspects of memory retrieval by modulating low-frequency oscillatory activity accordingly. However, it remains unknown whether and to what extent respiration might indeed coordinate retrieval-related power decreases in the α/β band and memory reactivation.

Here, we set out to directly assess the influence of respiration on the behavioral and neural expressions of memory retrieval using electroencephalography (EEG) and respiration recordings acquired during an episodic memory task. First, we replicate previous findings indicating that the phase of respiration at cue presentation influences memory performance, with successful retrieval being more likely when reminder cues were presented close to inhalation. We further show that this effect is potentially driven by respiration’s impact on both power modulations in the α/β band and concomitant memory reactivation. Moreover, the strength of the interplay between respiration and retrieval-related reactivation is directly associated with behavioral memory performance. Our results identify respiration as a potential pacemaker for episodic memory retrieval in humans and pave the way for future research on brain-body interactions during memory formation.

## Results

To examine whether respiration affects the behavioral and neural dynamics of memory retrieval, we analyzed EEG and respiratory recordings from 18 participants across two experimental sessions, each involving an episodic memory task (Fig. 1a). During the learning phase, participants were instructed to associate verbs with images of either objects or scenes, depending on the experimental session. Memory performance was evaluated in a sequential manner. Participants first completed a word-recognition task, classifying words (presented 3 s each) as “old” or “new.” For words recognized as “old”, participants attempted to retrieve the specific associated image (object or scene). Associative retrieval performance was normalized by the number of accurately recognized items (“hits”). Each session concluded with an independent “localizer task”, during which participants were presented with a new set of stimuli comprising both object and scene images. The data of the localizer task were used to train a linear classifier in order to distinguish object- and scene-related EEG patterns during memory retrieval. Note that parts of the data have already been published, showing that (i) slow oscillation-spindle complexes clock memory reactivation during NREM sleep^36^ and (ii) the occurrence of these sleep-related oscillations is modulated by respiration^4^.

**Figure 1.**
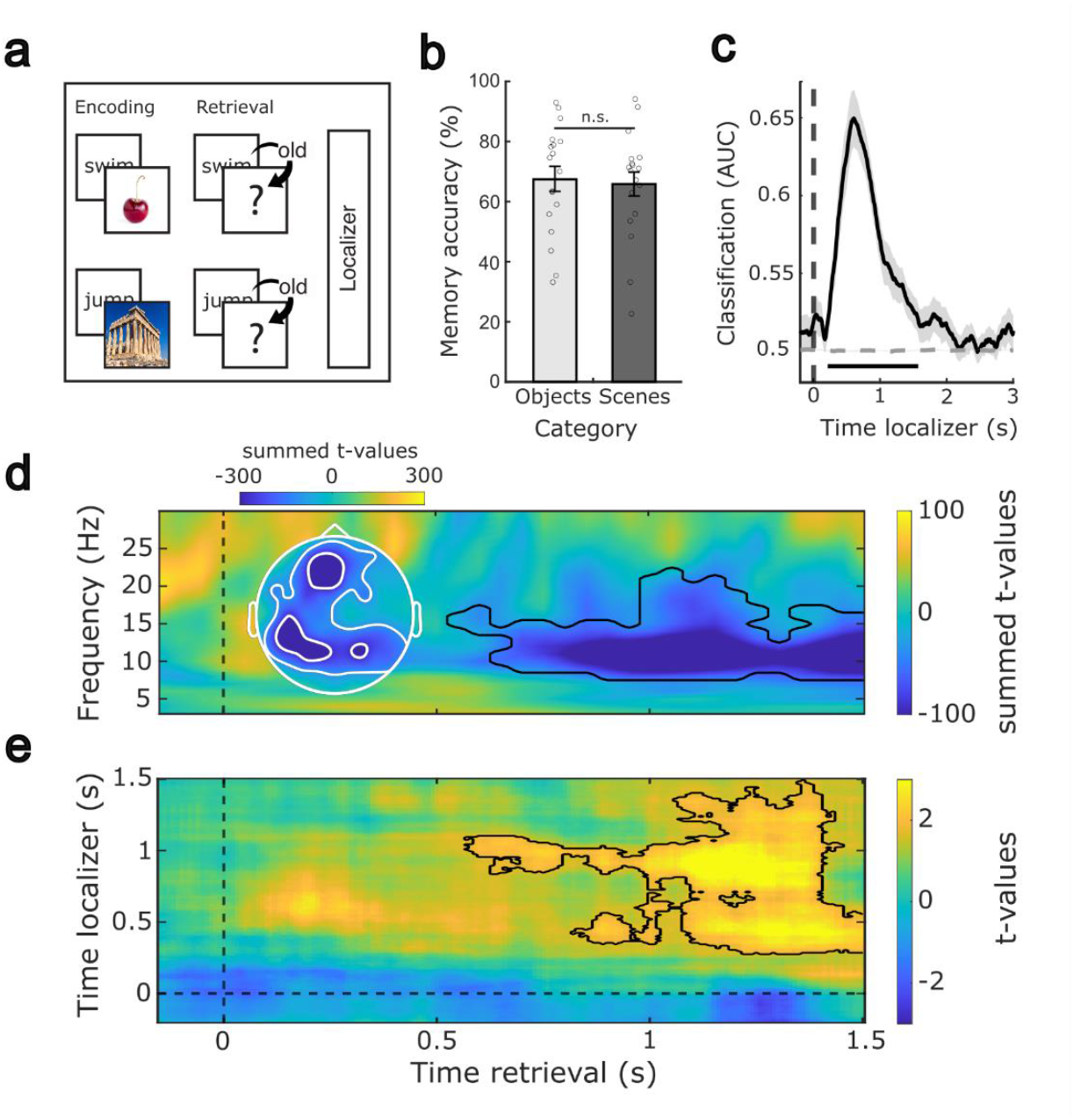
Experimental procedure, behavioral results, and retrieval-related EEG results. (a) During encoding, participants were presented with 120 verb-object or verb-scene pairs, depending on the experimental session, and memory performance was evaluated afterward. At the end of each session, participants performed a localizer task that involved processing a new set of object and scene images. (b) Behavioral results for both experimental sessions: objects (light gray) and scenes (dark gray). Bar graphs show the mean percentage (±SEM) of recalled image exemplars out of correctly recognized verbs. Dots indicate individual memory performance of participants (N = 18). No significant difference was found in memory accuracy between experimental sessions. (c) Stimulus categories (objects vs. scenes) could be decoded above chance from the localizer EEG data, starting around 150 ms post stimulus onset (the black solid line indicates decoding performance (±SEM)). The horizontal dashed line indicates surrogate decoding performance, which was estimated by shuffling the training labels 250 times. The vertical dashed line indicates stimulus onset (time = 0 s). The lower horizontal dark gray line shows the temporal extent of significant decoding results as derived from a dependent-samples t-test (one-sided, *p* < 0.001, corrected across time). (d) Remembered trials were associated with stronger power decreases in the α/β band (∼8-20 Hz) after cue onset (∼0.5 to 1.5 s) as compared to not remembered trials (contour lines indicate the extent of the significant cluster, *p* < 0.001, d = −1.5950; corrected across time, frequency and electrodes). The topographical inset illustrates the sum of t-values across all significant time points and frequencies for each electrode, illustrating that power differences were most prominent across frontal and parietal areas. (e) Learning-related brain patterns (objects vs. scenes) were decodable during retrieval (contour lines indicate the extent of the significant cluster, *p* = 0.032, d = 0.5972, one-sided; corrected across times); color range (blue to yellow) represents t-values comparing the decoding performance distributions of remembered versus not-remembered trials.

### Behavioral results and localizer-related classification

As previously reported^36^, memory performance was tested before and after a 120-minute nap (see Methods). To examine potential differences between the two learning categories (objects vs. scenes), memory scores were averaged across pre- and post-sleep sessions. No significant difference emerged between object and scene conditions (two-sided paired-sample t-test, t_(17)_ = 0.37, *p* = 0.71, d = 0.0872), suggesting that task difficulty was well matched across image categories (see Supplementary Table 1 for a detailed overview of memory measures). To derive the neural signatures of category-specific processing, which were used to track memory reactivation during retrieval, we employed a localizer task at the end of each session. During this task, participants viewed a novel set of images comprising both objects and scenes. Category-specific EEG patterns were extracted by aggregating the localizer data across experimental sessions and performing multivariate classification using linear discriminant analysis (LDA). Applying fivefold cross-validation, classification accuracy significantly exceeded chance levels between ∼150 ms and 1500 ms after stimulus onset (*p* < 0.001, corrected across time, Figure 1c), indicating that the classifier reliably captured category-specific neural patterns of object and scene image processing.

### Memory retrieval is associated with α/β power decreases and memory reactivation

As a first step, we set out to identify the oscillatory fingerprint of successful memory retrieval. To extract memory-related EEG patterns, we contrasted time-frequency representations between remembered and not-remembered items, based on associative memory test performance. In line with previous findings^24,29,37^, successful remembering was associated with significant power decreases in the α/β range (∼8 – 20 Hz), starting approximately 500 ms after cue onset across fronto-parietal areas (*p* < 0.001, corrected across time, frequency, and space; see Figure 1d). Next, we assessed whether neuronal activity related to the session-specific learning categories would be reactivated during successful retrieval^27,38^. Again, we trained an LDA classifier on the concatenated localizer data from both experimental sessions (objects and scenes; −0.15 to 1.5 s). The resulting training weights were then applied to retrieval data from both sessions, centered around cue onset (−0.15 to 1.5 s). The classifier testing labels matched the stimulus category from the preceding encoding session (object or scene), such that above-chance classification reflects activation patterns that more closely resemble the newly learned stimulus category than the alternative one. As shown in Figure 1e, results revealed significantly higher classification performance for remembered as compared to not-remembered items emerging between 0.5 and 1.5 s relative to cue onset (*p* = 0.032, corrected across times; see Supplementary Fig 1c for comparisons against chance-level distributions obtained via label shuffling).

### Respiration phase modulates memory performance and EEG activity during retrieval

To assess whether the phase of respiration (mean respiratory frequency = 0.267 ± 0.041 Hz) would impact memory performance during retrieval^20^, cue-centered trials (−0.5 to 3 s; see Supplementary Fig. 1 for the cue-centered respiratory event-related potential) were segmented into 250 ms bins. Each bin was then categorized according to its proximity to either an inhalation peak (i.e., the inhalation-to-exhalation transition) or an exhalation trough (i.e., the exhalation-to-inhalation transition). Memory accuracy for both categories was computed within each time bin and normalized by the participant’s individual overall performance. This yielded two time series per participant—one for bins aligned with the inhalation peak and one with the exhalation trough—capturing how changes in the respiratory phase over time impacted memory accuracy. A non-parametric cluster-based permutation test revealed that respiration significantly influenced retrieval performance (Fig. 2a, for recognition memory related results, see Supplementary Fig 1b). Specifically, successful retrieval was more likely in trials where the respiration cycle was closer to inhalation peaks between 0.125 and 0.625 s (*p* = 0.047, corrected across time) and to exhalation troughs between 1.875 and 2.375 s relative to cue onset (*p* = 0.044, corrected across time). Given the cyclical nature of respiration—moving from inhalation to exhalation—these findings suggest that trials in which participants inhaled around cue presentation and then transitioned to exhalation (within a trial) were more likely to be remembered.

**Figure 2.**
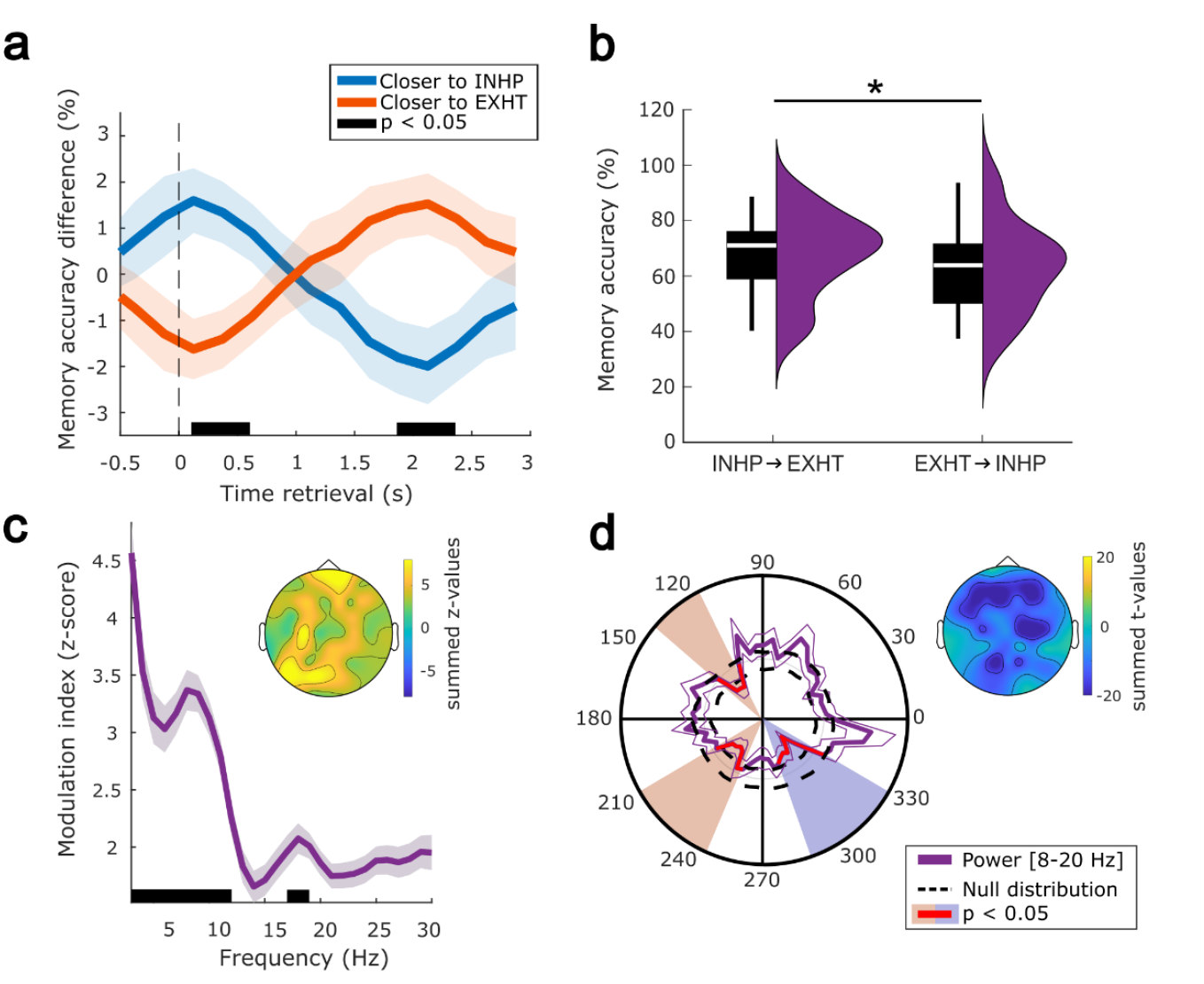
Respiration’s impact on memory performance and oscillatory EEG activity. (a) Successful retrieval was more likely in trials where participants inhaled during the period of cue presentation—with inhalation peaks occurring between 0.125 and 0.625 s (*p* = 0.047, d = 0.2927, corrected across time)—and subsequently exhaled — with exhalation troughs occurring between 1.875 and 2.375 s (*p* = 0.044, d = −0.3061, corrected across time). Blue / orange lines represent the difference in accuracy with respect to the participant’s overall memory performance for the trials that were closer to the inhalation peak /exhalation trough in each time bin (±SEM). The horizontal black line illustrates the temporal extent of significance as derived from a dependent-samples t-test (two-sided, cluster corrected across time). (b) Memory performance during retrieval was significantly enhanced when inhalation peaks preceded exhalation troughs (mean ± SD: 70.03±13.70) rather than the other way around (mean ± SD: 64.67 ± 17.11). Box plots show median (±Q_1/3_) accuracy of recalled image exemplars out of correctly recognized verbs. Star denotes a significant difference as derived from a two-sided, dependent-samples t-test (t_(17)_ = 2.868, *p* = 0.011, d = 0.6761). (c) The modulation index (± SEM, in shade grey) suggests that the phase of respiration significantly influences EEG amplitudes in the α/β range with local peaks at 9 Hz (z > 2.58) and 18 Hz (z > 1.96). The lower horizontal black line depicts all significantly modulated frequencies (*p* < 0.05, corrected across frequencies using FDR). (d) Power in the α/β band is modulated at different phases of the respiratory cycle (purple solid line represents the power of [8-20 Hz] band (±SEM)). Segments in red indicate phase bins associated with significant power decreases in the α/β range as derived from a dependent-samples t-test against a shuffled reference distribution (114 to 138°, p = 0.023, d = −1.4681; 210 to 246°, p = 0.043, d = −1.3003; 288 to 330°, p = 0.003, d = −1.4326; two-sided, cluster corrected across phase and electrodes; inhalation peak = 0°). Light blue (orange)-shaded areas represent significant phase bins closer to the inhalation peak (exhalation trough). The topographical insert illustrates the sum of t-values across all significant phases for each electrode, showing that this decrease was more prominent in frontal, parietal, and right-central cortices.

To further corroborate this assumption, we divided the trials into two groups: one where inhalation peaks occurred before exhalation troughs during retrieval time and another where the order was reversed. Next, memory accuracy was computed for each of the categories. As shown in Fig. 2b, memory performance during retrieval was significantly enhanced when inhalation peaks preceded exhalation troughs rather than the other way around (dependent samples t-test, two-sided, t_(17)_ = 2. 868, *p* = 0.011). Thus, retrieval was more likely when cues occurred prior to inhalation peaks, with an additional increase in retrieval probability around exhalation troughs later in the trial. The follow-up sequence analysis clarified that these effects reflect a complementary pattern: memory performance was highest when an inhalation at cue onset was followed by exhalation thereafter.

To examine whether respiration would likewise modulate EEG activity during retrieval, we calculated the modulation index (MI)^39^ to estimate cross-frequency phase-amplitude coupling between respiration (providing the phase) and EEG signals across all electrodes (providing amplitude measures between 3–30 Hz). For statistical assessment, we generated an MI-level reference distribution by randomly shifting the phase-related time series of each trial 200 times. The mean and standard deviation across the reference distribution were then used to z-score the MI of the empirical distribution. Results revealed a significant modulation of EEG amplitudes by the phase of respiration with local peaks in the α (9 Hz, z > 2.58 / *p* < 0.01) and β range (18 Hz, z > 1.96 / *p* < 0.05; Fig. 2c). Next, to specifically characterize EEG fluctuations in the α/β band across the respiratory cycle, we divided the respiratory phase of each trial into 60 evenly spaced bins (bin size = 6°). Then, the mean power in the α/β band (i.e., 8–20 Hz) was computed for each phase bin across all trials and contrasted against a null distribution (see Methods). The 8–20 Hz range was chosen because it corresponds to the α/β band that showed the most consistent retrieval-related power decreases in our dataset (Fig. 1d) and in previous studies^25,38,40^. This predefined band was therefore used to examine respiration-locked modulation of retrieval-relevant oscillatory activity. We found respiration-locked EEG power exhibited significant decreases around the exhalation through (114 to 138°, *p* = 0.023; 210 to 246°, *p* = 0.043) and towards the inhalation peak (288 to 330°, *p* = 0.003; all corrected for multiple comparisons across phase bins and electrodes, see Fig. 2d).

### Respiration modulates retrieval-related memory reactivation and its interplay with α/β power

Having established that respiration modulates both retrieval-related memory performance and EEG activity in the α/β range during retrieval, we finally asked whether respiration would also impact the emergence of memory reactivation. To this end, we detected exhalation troughs, the most prominent feature of the cue-locked respiratory ERP (see Supplementary Fig. 1a), in the respiratory signal during retrieval (0 to 3 s relative to cue onset) and centered the EEG data accordingly. Again, a classifier was trained on the pooled localizer data from both learning sessions (− 0.15 to 1.5 s), but the resulting training weights were this time specifically applied to the exhalation trough-locked EEG segments (±2 s, accounting for a whole respiratory cycle with a mean respiratory frequency of 0.25 Hz). Results revealed significantly higher decoding performance for remembered as compared to not-remembered trials emerging just prior to the exhalation through (p = 0.042, corrected across times, Fig. 3a; see Supplementary Fig 2a for remembered and not remembered against chance-level distributions obtained via label shuffling). However, while these effects support the notion of retrieval-related reactivation, they should nonetheless be interpreted with caution given their moderate size.

**Figure 3.**
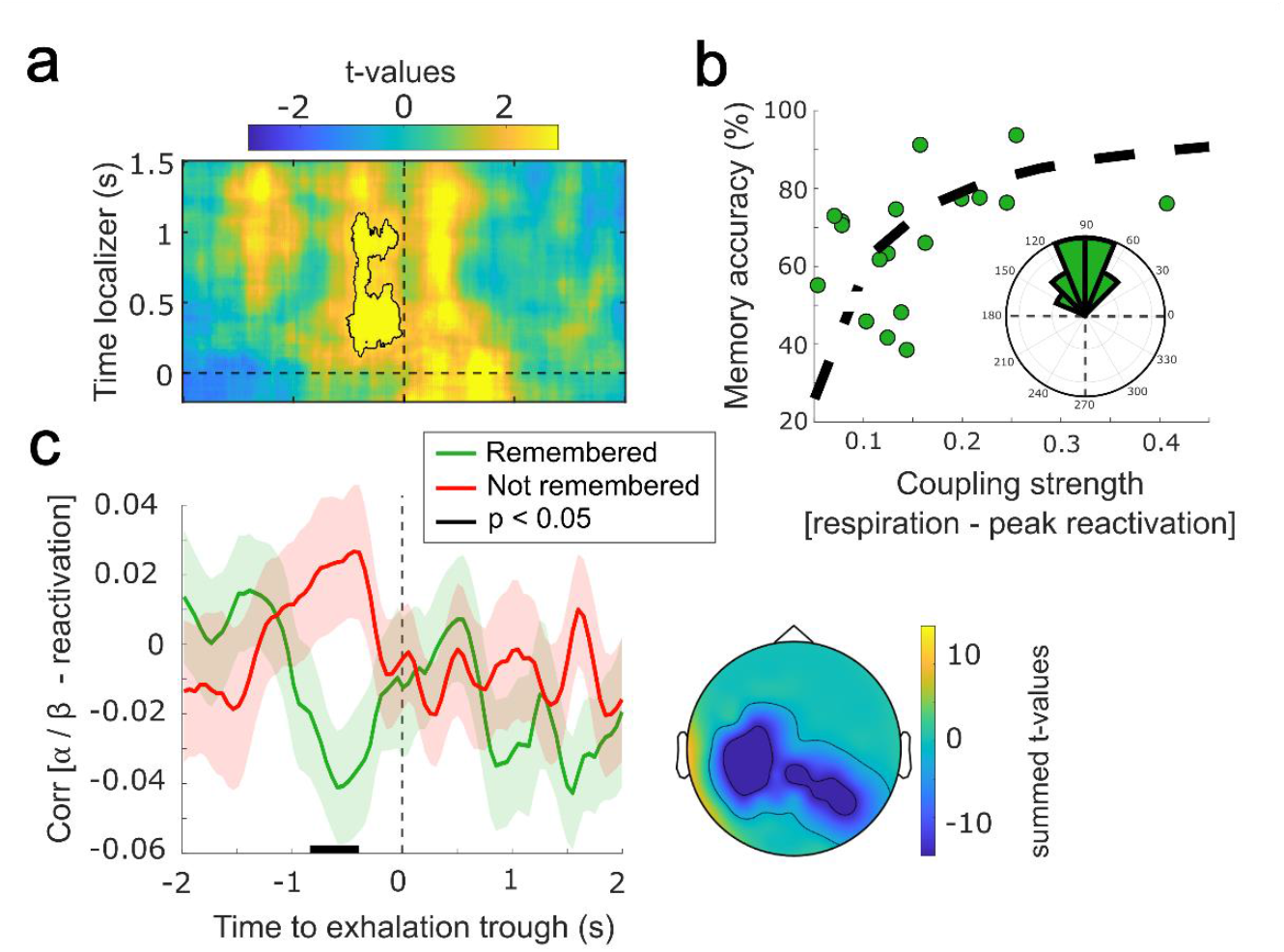
Respiration impacts memory reactivation and concomitant α/β power fluctuations. (a) Learning-related brain patterns (objects vs. scenes) were decodable during retrieval based on exhalation trough-centered EEG data. Contour lines indicate the extent of the significant cluster (*p* = 0.042, d = 0.8434, one-sided, corrected across times); color range (blue to yellow) represents t-values of remembered against not-remembered distributions of decoding performances. (b) In remembered trials, coupling strength (measured as vector length) between respiration and classification maxima is positively correlated with memory performance (Spearman’s Rank Correlation, r = 0.558, *p* = 0.018). Nonlinear fit as a dashed black line (only for visualization purposes). The polar histogram illustrates the mean respiratory phase at which decoding-derived reactivation peaks emerge during remembered trials for each participant (mean angle = 93.045º ± 0.321; Rayleigh test z = 14.58, *p* < 0.001; inhalation peak = 0°). (c) Decoding-derived single trial decision values and α/β power (8-20 Hz) are more negatively correlated prior to the exhalation trough in remembered trials as compared to not-remembered trials. The green (red) solid line shows the Spearman’s correlation for (not) remembered trials (±SEM). The horizontal black line shows the temporal extent of significant results as derived from a dependent-samples t-test (two-sided, *p* = 0.023, d = −1.0248, cluster corrected across time and electrodes). The topographical inset illustrates the sum of t-values across all significant timepoints for each electrode, showing that this difference was stronger in the parietal cortex.

To further assess the functional significance of the coupling between memory reactivation and respiration, we determined its impact on the participants’ memory performance. To this end, we extracted decision values from the decoding procedure, yielding a single-trial, time-resolved metric that reflects the classifier’s confidence in assigning a specific category (objects vs. scenes) in the cue-centered EEG data^41^. Then, for each of the remembered trials, we extracted the respiratory phase where decision values reached their maximum. We found that decision value maxima consistently emerged during exhalation across participants (mean angle = 93.045º ± 0.321; Rayleigh test z = 14.58, *p* < 0.001; see polar insert in Fig. 3b). Finally, when correlating the vector length (representing the coupling strength between respiration and classification maxima across trials) with memory performance, we observed a significant positive relationship (Fig. 3b; Spearman’s Rank Correlation, r = 0.558, *p* = 0.018; for results based on non-remembered trials see Supplementary Fig. 2). The distribution indicated that the stronger the reactivation-related EEG signatures were modulated by respiration, the better the memory performance.

So far, we have demonstrated that respiration might serve as a temporal anchor for retrieval-related neural processes, as both characteristic power decreases in the α/β band and memory reactivation were significantly modulated by the phase of respiration. Hence, we hypothesized that respiration might co-modulate both key markers of successful memory retrieval. Thus, we extracted trial-based decision values, locked to exhalation troughs, and correlated them with α/β power values at each electrode and timepoint across trials. Again, we focused on the predefined 8–20 Hz α/β range identified as most consistently linked to retrieval-related power decreases (Fig. 1d), ensuring that respiration–reactivation coupling was assessed within the same functionally relevant frequency band. When contrasting correlational values between remembered and not-remembered trials, we observed a significant negative cluster emerging just prior to the exhalation through (*p* = 0.023; corrected for multiple comparisons across time and electrodes, Fig. 3c), indicating that the stronger the respiration locked α/β power decrease, the higher the classifiers fidelity during successful remembering.

## Discussion

Our results unveil a role of respiration in modulating both behavioral and neural components of remembering. In line with previous research, we show that the phase of respiration influences memory performance (Fig. 2a), with successful retrieval being more likely when reminder cues were presented close to inhalation. Moreover, key neural signatures of retrieval, oscillatory power decreases in α/β band (Fig. 2d) and memory reactivation (Fig. 3a), as well as their interplay (Fig. 3c) were modulated by respiration. Finally, the strength of the coupling between respiration and reactivation processes was associated with memory performance (Fig. 2b), underscoring the functional relevance of respiration–brain interactions for memory.

Extensive research in both rodents^3,11,42^ and humans^2,6,20^ indicates that respiration might be essential for modulating neural oscillations across various brain regions and frequency bands. Mechanistically, it has been suggested that the impact of respiration on neural activity is mediated by incoming airflow stimulating mechanoreceptors in olfactory sensory neurons^12^. Through this stimulation, breathing-locked oscillations are generated and then relayed to the olfactory cortex and spread to various target regions, and in particular to sensory areas^9,10,12^. These oscillations transiently increase neural excitability and may thus facilitate stimulus perception^16^. Beyond sensory processing, respiration has also been shown to impact cognition in various domains such as motor control^17,18^ and emotion^21^, often enhancing performance when stimuli are presented during inhalation^7^. In particular, a growing number of findings in humans demonstrate that breathing impacts memory processes during wake^23 (but see 43,44)^. Zelano and colleagues were the first to show that during retrieval testing, memory was better for stimuli that were presented during inhalation as compared to exhalation^20^. Interestingly, this effect became only evident when participants engaged in nasal breathing (as compared to mouth breathing), suggesting a specific role for nasal respiration in orchestrating memory-related processes. Our data remain agnostic as to whether nasal breathing is indeed crucial for impacting memory performance. Nevertheless, our finding that successful retrieval was more likely when participants inhaled during reminder cue presentation is in line with the previous result, indicating that the phase of respiration at the time of cue presentation impacts memory performance. However, while these results appear to suggest that stimulus processing might be facilitated during inhalation, the neural mechanisms underlying respiration’s influence on successful memory retrieval have remained unexplored. Notably, another study examining respiration and memory^44^ did not observe performance differences across the respiratory cycle. This discrepancy may relate to task design: Schaefer et al. used a recognition task without associative retrieval demands, whereas our paradigm required retrieval of specific verb–image associations. Indeed, when we restricted our analysis to recognition trials, no significant modulation by respiration phase was found (Supplementary Fig. 1b). Taken together, these observations suggest that respiration effects may emerge more prominently in tasks requiring associative retrieval or stronger mnemonic reconstruction than in simple recognition paradigms.

Cortical power decreases in the α/β band have consistently been reported to index successful memory retrieval^24–27^. It has been suggested that these decreases might facilitate remembering by allowing neural assemblies to express the specific neural codes associated with stored memories^25 (but see 45)^. Indeed, α/β power modulations have been shown to track the amount and content of information being retrieved^25,46,47^, the fidelity of memory details^28–30^, and potentially provide time-windows of opportunity for memory reactivation^24,35,48^. In our data, successful retrieval, irrespective of respiration, was associated with robust power decreases in the α/β band across frontal and parietal areas. Two separate learning sessions with distinct stimulus sets (objects vs. scenes; Fig. 1a) enabled us to track retrieval-related reactivation and demonstrate that learning-specific patterns were decodable during periods of low α/β power.

Beyond these confirmatory findings concerning the role of low-frequency power decreases and concomitant memory reactivation in memory retrieval, our results suggest that respiration might play a crucial role in modulating those neural dynamics underlying successful remembering. Fluctuations in α/β activity have already been demonstrated to be influenced by respiration during diverse cognitive tasks (i.e., visuo-spatial, lexical, and math)^19^, perception^16^, and rest^6^. Our results critically extend these findings by demonstrating that also retrieval-related α/β fluctuations are tightly modulated by respiration. Specifically, we found that decreases in α/β activity systematically emerged both around exhalation troughs and just prior to inhalation peaks. These power reductions, synchronized with different phases of the respiratory cycle, may serve distinct functional roles. As outlined above, our behavioral analyses revealed that successful remembering was more likely when participants inhaled during the reminder cue presentation and exhaled thereafter (i.e., with exhalation troughs occurring between 1.875 and 2.375 s after cue onset). Interestingly, in addition to their role in memory retrieval, low pre-stimulus α/β levels have consistently been linked to enhanced perceptual performance^49^. Thus, inhalation-locked decreases in low-frequency power may facilitate stimulus perception^16^, forming a fundamental foundation for cue-triggered remembering. On the contrary, signatures of retrieval-related memory reactivation are typically observed between ∼500 and 1500 ms after reminder cue onset^38^. Accordingly, low-frequency power decreases emerging around exhalation troughs might establish a neural environment that is optimally tuned for memory reactivation to unfold. This interpretation is in line with our finding that not only power decreases in the α/β band were modulated by exhalation, but that classification-derived memory reactivation, hence the putative neural substrate of remembering^31^, emerged just prior to the detected exhalation troughs. Therefore, our results add further evidence to the notion that that different phases of respiration might establish time windows of opportunity for distinct brain functions^9^. In addition, respiration has recently been established to similarly influence the emergence of sleep-related oscillations and in consequence, memory reactivation in humans^4^, with reactivation processes putatively unfolding during exhalation. Together, these results suggest that respiration might impact memory-related neural oscillations and concomitant memory reactivation in a consistent manner, regardless of state. One potential concern of our paradigm, however, was that participants were required to withhold their responses for three seconds, ensuring that the respiration effects we report occurred well before any motor output. We therefore interpret the observed exhalation-locked neural dynamics as reflecting retrieval rather than response execution. Nevertheless, future work without a response delay will be valuable to further disentangle how respiration may jointly coordinate internal retrieval and overt responding.

The negative correlation between α/β power and memory reactivation signatures prior to exhalation troughs suggests that respiration may shape the interaction between two key neural markers of successful remembering. However, it remains unclear whether respiration directly shapes the emergence of memory reactivation or acts indirectly via its effects on α/β activity. Recent studies suggested that α/β power decreases may reflect eye movements rather than memory processes per se^45,50^. In this light, our findings fit within a broader brain–body framework, in which respiration not only modulates cortical excitability^16,51^, but may also influence, or be influenced by, oculomotor behavior^52,53^, thereby contributing to fluctuations in α/β power. To disentangle these intertwined processes, future studies combining respiration measures, eye tracking, and neural recordings will be essential. Moreover, our design is correlational and cannot determine whether respiration actively drives retrieval-related neural dynamics or whether both are co-modulated by a third factor (e.g., arousal, attention, or autonomic tone). Causal manipulations of breathing will therefore be crucial.

Complementary intracranial recordings could further test whether respiration modulates the occurrence of hippocampal ripples during retrieval^54^, which are known to trigger memory reactivation in both hippocampal and cortical regions^55^. Given that ripple occurrence has been shown to be modulated by respiration in rodents^3,56^ and more recently in humans^57^, particularly emerging during exhalation, such work may clarify whether exhalation provides a temporal window of reduced neural noise that facilitates mnemonic reactivation. In any case, our finding that the strength of respiration–reactivation coupling predicts individual memory performance provides compelling evidence for the functional relevance of this relationship, irrespective of whether respiration exerts a direct or indirect influence on reactivation processes. Nevertheless, while our analyses were specifically locked to the respiratory cycle, and the observed effects align with prior reports of respiration–brain coupling^6,20^, we cannot exclude the possibility that other physiological rhythms (e.g., cardiac cycle, slow cortical potentials) contribute to the observed modulations^58^. Future studies concurrently measuring respiration, cardiac activity, and other slow fluctuations will be essential to disentangle their respective influences on retrieval‐related neural dynamics.

To conclude, our results reveal a critical role for respiration in modulating both the behavioral and neural aspects of remembering. They identify respiration as a potential pacemaker for effective remembering and highlight the importance of brain-body interactions in human memory processing.

## Methods

The current analyses are based on reference^36^, where detailed information about participants, stimuli, task, and data acquisition can be found. The data of two participants (out of 20) had to be excluded due to low trial numbers (< 15 trials in any of the conditions). In summary, the study involved 18 healthy participants (mean age: 20.89 ± 0.34; 15 female). Pre-study screening, which included the Pittsburgh Sleep Quality Index (PSQI^59^), the morningness-eveningness questionnaire^60^, and a custom questionnaire assessing general health and stimulant use, confirmed that participants were not taking any medication and had no neurological or psychiatric conditions at the time of the experiment. The University of Birmingham Research Ethics Committee approved the study, and all participants provided written informed consent. They were compensated either financially or with course credit for their participation.

### Experiment overview

The experiment involved two sessions per participant, spaced at least one week apart (mean interval: 7.78 ± 0.43 days), with the session order counterbalanced across participants. On each experimental day, participants arrived at the sleep laboratory at 11 a.m., where the set-up for polysomnographic recordings began. Electrodes for electroencephalography (EEG), electromyography (EMG), and electrocardiography (ECG) were applied, along with a thermistor airflow sensor to monitor breathing. Around noon, the experiment commenced with a modified psychomotor vigilance task (PVT^61^), followed by a memory task (see Encoding and Memory tests below) in which participants learned to associate 120 verbs with images (either objects or scenes^62^, depending on the session). The sleep period began at approximately 1 p.m., giving participants the opportunity to nap for ∼120 minutes. After the nap, their vigilance was reassessed using the PVT, and memory performance was tested again. A localizer task was conducted at the end of each session (see Localizer task below). Behavioral responses and the presentation of tasks were managed using Psychophysics Toolbox Version 3^63^ and MATLAB 2018b (MathWorks, Natick, USA).

#### Stimuli

The experimental stimuli consisted of 360 verbs and 240 images, evenly divided between object and scene categories, presented during both experimental sessions. Object images depicted animals, food items, clothing, tools, or household items against a plain white background (e.g., a calculator). Scene images represented identifiable landscapes or locations (e.g., an island). These categories were chosen as they engage distinct brain networks, such as the lateral occipital complex for object processing^64^ and the para-hippocampal place area for scene processing^65^, facilitating the investigation of experience-driven memory reactivation during retrieval (see Methods: Multivariate analysis). All images were obtained from reference ^62^.

#### Familiarization

The experiment commenced with an image familiarization phase designed to (i) facilitate the learning of verb-image associations during the subsequent encoding session and (ii) establish accurate image labels for later cued recall. Each trial began with the presentation of a fixation cross for 1.5 ± 0.1 s. Participants were then shown one of 130 images depicting either objects or scenes, depending on the experimental session. Of these, 120 images were included in the learning material and accompanied by captions accurately naming the depicted exemplars. An additional 10 images, which were not used further in the experiment, were paired with incorrect captions. Each stimulus was displayed for 2.5 s. Participants were instructed to press a button upon detecting an erroneous image-caption pairing.

#### Encoding

Each trial began with the presentation of a fixation cross for 1.5 ± 0.1 s, followed by a verb (e.g., “jump”) displayed for 1 s. Immediately after, the corresponding image to be associated with the verb was presented for 4 s. Participants were instructed to create a vivid mental image or narrative linking the verb with the object or scene depicted in the image. Following the image presentation, participants indicated whether the mental image they had formed was realistic or bizarre. Participants were informed in advance that their memory for the verb-image pairs would be tested subsequently. The learning block was repeated twice with a randomized trial order in each iteration to achieve satisfactory memory performance, as established in a prior pilot study.

#### Memory tests

Participants’ memory for the encoded verb-image pairs was assessed both before and after the nap interval. To mitigate potential testing effects on behavioral measures of memory consolidation^66,67^, only half of the learned verb-image pairs were included in each memory task, with no overlap between the two tests. Each memory test comprised 60 randomly selected verbs from the learning phase, intermixed with 30 novel verbs (“foils”) that had not been presented during learning. Each trial began with a fixation cross displayed for 1.5 ± 0.1 s, followed by the presentation of a verb on the screen. After 3 s, participants indicated within a 10 s window whether the verb was “old” (i.e., part of the learning material) or “new” (i.e., not seen during learning). For “new” responses, participants proceeded directly to the next trial. For “old” responses, participants were prompted to either type a description of the associated image or enter “do not know” if they could not recall the target image. Responses were scored as correct if (i) the typed description matched the original caption presented during the familiarization phase or (ii) the description unambiguously corresponded to the content of the target image.

#### Localizer task

During the localizer task, participants viewed a new set of images comprising 90 objects and 90 scenes, irrespective of the experimental session. Each trial began with a fixation cross presented for 1.5 ± 0.1 s, followed by the display of a randomly selected image (object or scene) on the computer screen for a duration between 2.5 and 10 s. Each image was presented twice during the task, and participants were instructed to indicate whether the image was being shown for the first time (“new”) or the second time (“old”).

### EEG and respiration

A Brain Products 64-channel EEG system was used to record electroencephalography (EEG) throughout the experiment. Impedances were kept below 10 kΩ. EEG signals were referenced online to electrode FCz and sampled at a rate of 1000 Hz. Respiration was recorded using an Embla thermistor airflow sensor.

### Data analysis

#### Behavioral

Analysis was performed based on associative memory performance, calculated as the proportion of correctly remembered images relative to the number of ‘hit’ trials (i.e., verbs correctly identified as old), separately for each session (object and scene) and memory test (pre- and post-sleep). To relate the respiratory phase to memory performance across trials (Fig. 2a), we first collapsed all trials from both sessions and memory tests (i.e., pre- and post-sleep). Then, trials were segmented into 250 ms bins, starting −0.5 s before cue onset. For each time bin, we categorized trials based on whether the circular mean of the respiration phase across the 250 ms of a bin aligned more closely with the inhalation peak or the exhalation trough, respectively. Memory accuracy was then computed for each category (inhalation and exhalation relatedness, respectively) within each time bin and normalized by each participant’s overall memory performance. Finally, to analyze whether specific respiration sequences (inhalation to exhalation or exhalation to inhalation) during retrieval influence memory performance, we categorized trials based on the presence of either an inhalation peak followed by an exhalation trough or the reverse within −0.5 to 3 s from cue onset. Then, associative memory performance was computed for each category.

#### EEG and respiration

All data were analyzed using MATLAB (2020a; MathWorks). EEG data were preprocessed using the FieldTrip toolbox^68^ for EEG/MEG analysis (v.16/07/2023). All data were downsampled to 200 Hz and re-referenced using a common average reference (CAR). Noisy EEG channels were identified by visual inspection, discarded, and then interpolated using a weighted average of the neighboring channels. EEG epochs related to the retrieval task were locked to two different events: stimulus presentation and exhalation troughs. Exhalation troughs (and inhalation peaks for Fig. 2b) were detected using the Breathmetrics toolbox^69^. After detecting exhalation troughs and inhalation peaks from the respiratory signal, trials were classified as INHP → EXHT when an inhalation peak preceded an exhalation trough within −0.5 to 3 s around cue onset (i.e., the same time window used in Fig. 2a), and as EXHT → INHP for the reverse order. This classification was based on detected respiratory events rather than phase estimates, minimizing potential bias from phase interpolation. In addition, trials that contained an exhalation trough within 3 s after cue onset were aligned to that respiratory event for analyses shown in Fig. 3, as exhalation troughs constituted the most robust and temporally consistent respiratory marker across trials (Supplementary Fig. 1). Trials were classified into remembered and not-remembered categories based on associative memory performance. In the case of localizer trials, all data were locked to the stimulus onset with a temporal window of −0.2 to 3 s. For analyses related to respiratory phase, the phase was extracted using the double interpolation method^4,16^, unless specified otherwise.

#### Time-frequency analysis

Time-frequency analysis of stimuli-locked (−2 to 3 s) and respiratory event-centered EEG segments (−5 to 5 s) was performed using FieldTrip^68^. The longer time segments were chosen to allow for resolving low-frequency activity within the time windows of interest for both stimuli-locked (−0.2 to 1.5 s) and respiratory event-locked (−2 to 2 s) and avoid edge artifacts. Frequency decomposition of the data was achieved using Fourier analysis based on sliding time windows (moving forward in 50 ms increments). The window length was set to five cycles of a given frequency (frequency range: 1–30 Hz in 1 Hz steps). The windowed data segments were multiplied with a Hanning taper before Fourier analysis. Afterward, power values were z-scored across time, using a 1 s baseline window prior to stimulus onset. To analyze oscillatory power as a function of the respiratory phase, power values were transformed from the time axis to the phase axis. For that, the respiratory cycle was divided into 60 non-overlapping phase bins (bin size = 6°), and after assigning all power values to the respiratory phase bin where they occurred, the mean power was computed for each bin^16,51^. Thus, we obtained a respiratory phase-resolved distribution, with the value at each phase bin representing the mean of all power values that overlapped with that respiratory frequency across all trials from 0 to 3 s.

#### Modulation Index

Phase amplitude coupling was assessed using the Modulation Index (MI)^39^. To estimate the instantaneous phase of the respiration signal, we filtered the continuous respiratory data around the peak frequency of each particular subject ± 0.05 Hz two-pass Butterworth band-pass filter). We then extracted instantaneous amplitude data across frequencies between 3 and 30 Hz at each electrode in steps of 2 Hz regardless of the α/β regions identified in Fig. 1d, to avoid circularity and because respiration has been shown to modulate large-scale neural networks. To this end, a two-pass FIR filter (order = three times the lower frequency bound) was used to create 27 equally spaced frequency bins, with center frequencies ranging from 3 to 30 Hz, and with fixed-frequency bandwidths of 2 Hz. The envelope of the Hilbert-transformed band-pass filtered data was then used as an amplitude estimate. To compute the MI (for a given frequency pair), we divided the phase signal into 20 bins, and then, for each bin, computed the mean amplitude. This yields a distribution of amplitude as a function of phase. The MI is defined as the Kullback–Leibler distance between that distribution and the uniform distribution (across the same number of bins). To assess the statistical significance of MI values, we generated participant-specific null distributions by randomly shifting the respiration phase time series relative to the neural signal (200 iterations per participant). The mean and standard deviation of each null distribution were used to z-score the empirical MI values, which were then converted into p-values. This phase-shuffling procedure preserves individual respiratory dynamics while providing within-subject reference estimates. Additionally, we equalized the number of correct and incorrect trials within each participant via subsampling. Results were corrected for multiple comparisons within each channel using FDR across frequencies. The channels with frequencies that survived FDR correction were then averaged, and frequencies laying over z > 1.96 (i.e., p < 0.05) were considered significant.

#### Multivariate analysis

Multivariate classification of single-trial EEG data was performed using MVPA-Light, a MATLAB-based toolbox for multivariate pattern analysis^70^. For multivariate classification, Linear Discriminant Analysis (LDA) was used as classifier^70^. Prior to analyses the raw EEG data were bandpass-filtered between 0.1 and 45 Hz (4^th^ order Butterworth filter) and then z-scored the localizer data across all trials independently for each time point. Data from both sessions were then combined and subjected to principal component analysis (PCA), which transforms the data into linearly uncorrelated components ranked by the variance explained by each component^71^. PCA was employed to reduce dimensionality and mitigate overfitting, retaining the first principal components that explained 95% of the data’s variance^72^. To assess whether object and scene representations could be distinguished in the localizer, we trained and tested a classifier to discriminate between object and scene trials. A 200 ms running average smoothing window was applied to the data. EEG channels were treated as features, and a separate classifier was trained and tested at each time point. Classification performance was quantified using the area under the receiver operating characteristic curve (AUC), which measures the probability that a randomly selected pair of Class A and Class B trials is correctly assigned to their respective classes (0.5 = chance level; 1.0 = perfect classification). To prevent overfitting, fivefold cross-validation was implemented, with the analysis repeated five times to account for variability arising from random fold assignments^73^. The results were averaged across iterations. Statistical significance was evaluated by generating surrogate decoding performance values through 250 iterations of label shuffling, producing baseline values under the null hypothesis of label exchangeability. Classification performance was evaluated against this null distribution with a non-parametric cluster-based permutation test (see Statistics). This test was performed one-sided, as the relevant hypothesis was strictly directional: we expected stronger decodability of the current visualized category within the localizer trials.

To investigate differential evidence for object vs. scene representations as a function of prior learning during the retrieval task, we used the temporal generalization method^74^. First, we applied the same preprocessing described above to the concatenated localizer and retrieval data. A classifier was trained at each time point in the localizer data and applied to each time point during retrieval, specifically for remembered and not-remembered trials. No cross-validation was required since localizer and retrieval datasets were independent. As a metric, we again used the AUC. Remembered and not-remembered distributions were tested against each other again under a unidirectional hypothesis, as based on prior work, we expected stronger reactivation of the originally learned category during remembered compared to not-remembered trials. We additionally tested classifier performance against label-shuffled surrogate distributions (200 permutations) for remembered and not-remembered trials, separately for cue- and exhalation-locked retrieval data. The surrogate data underwent identical preprocessing, with PCA components retained up to 99 % of explained variance (instead of 95 % in the main analysis) to ensure numerical stability of the classifier during repeated training on label-shuffled data. Decoding results were then compared with their respective null distributions to identify significant above-chance time windows^41,75^ (see Supplementary Figs. 1c, 2a).

For trial-based analyses (Fig. 3b-c), decision values^41^ were extracted at each time point in the localizer and retrieval phases for each individual trial of the retrieval task (cue-centered for Fig. 3b; exhalation trough-centered for Fig. 3c). These values were then averaged across the significant time window in the localizer (150 to 1500 ms), resulting in a trial-by-retrieval time matrix. The modulus of the decision values was used to represent the model’s confidence at each time point of every trial, regardless of the trial’s original category. For the analysis in Fig. 3b, the maximum decision value of each trial was calculated to obtain the respiratory phase at peak reactivation.

#### Statistics

Behavioral retrieval data were averaged across conditions (object- and scene-condition, pre- and post-sleep) and subjected to a two-sided dependent-sample t-test (Fig. 1b for category comparison). To examine the effect of the sequence order of respiratory events (Fig. 2b) on memory performance, a two-tailed paired-sample t-test was conducted. For correlation analyses, Spearman’s correlation was used. To assess the relationship between oscillatory power (8 to 20 Hz) and memory reactivation (decision values) across exhalation trough-locked retrieval times, correlational values were smoothed using a 250 ms moving mean before statistical analysis. Effect sizes from statistical tests were computed by Cohen’s d.

All data were collapsed across both sessions (objects and scenes) and memory tasks (pre- and post-sleep) per participant before entering statistical analysis. Unless stated otherwise, we used non-parametric cluster-based permutation tests implemented in FieldTrip^68^, which do not rely on the assumption of a specific underlying distribution. A dependent-samples t-test was conducted at the sample level to identify clusters of contiguous time points across participants, with values thresholded at *p* = 0.05. The cluster statistic was defined as the maxsum (sum of all t-values within a cluster), and Monte Carlo simulations were performed to determine the cluster *p*-value (alpha = 0.05, two-tailed, unless stated otherwise) based on the permutation distribution. For circular analyses, data were divided into 60 distinct phase bins (bin length = 6°), and mean values were calculated within each bin. To assess statistical significance, the time course values within each trial were shuffled across trials 1000 times per subject and electrode, maintaining the original respiration phase to create a null distribution for each participant. This conservative approach ensures that the temporal characteristics of the respiratory signal is preserved^76^. Significance was determined by non-parametric cluster-based permutation tests for circular data.

## Data availability

All data supporting the findings of this study are publicly available at Open Data LMU (https://data.ub.uni-muenchen.de/596).

## Code availability

All code related to the analyses of the manuscript is available at GitHub (https://github.com/EstebanBT/Bullon-Tarraso-2025)

## Acknowledgements

T. Sc. is supported by the Emmy Noether program of the German Research Foundation (492835154).

## Contributions

T.Sc., M.P. and B.P.S. conceived the study and designed the experiment. T.Sc. conducted the experiment. E.B.T and T.Sc. analyzed the data. E.B.T., F.S, M.P., T.St., B.P.S. and T. Sc. wrote the paper.

## Supplemental Information

**Supplementary Table 1.** Overview of memory performance. Associative memory % refers to the percentage of correctly recalled images (relative to the total number of stimuli), while associative memory [out of hits] refers to the percentage of recalled image exemplars out of correctly recognized verbs. Statistical differences between conditions (objects vs. scenes) were assessed using dependent samples t-tests (two-sided).

|  | Objects | Scenes | *t* | *p* |
| --- | --- | --- | --- | --- |
| Recognition [Hits] % |  |  |  |  |
| pre-sleep | 74.14±3.97 | 73.80±4.15 | 0.12 | 0.90 |
| post-sleep | 64.54±3.98 | 66.05±4.45 | -0.60 | 0.56 |
| post- relative to pre-sleep | 87.31±2.85 | 88.68±2.34 | -0.39 | 0.70 |
| Recognition [Correct Rejections] % |  |  |  |  |
| pre-sleep | 90.00±2.40 | 87.41±4.70 | 0.60 | 0.56 |
| post-sleep | 88.33±3.19 | 84.63±5.70 | 0.72 | 0.48 |
| post- relative to pre-sleep | 97.77±1.72 | 96.22±3.92 | 0.43 | 0.67 |
| Associative Memory % |  |  |  |  |
| pre-sleep | 52.77±4.71 | 51.36±4.67 | 0.40 | 0.70 |
| post-sleep | 42.75±4.88 | 39.08±3.84 | 0.87 | 0.40 |
| post- relative to pre-sleep | 79.28±4.83 | 76.43±4.05 | 0.38 | 0.71 |
| Associative Memory [out of hits] % |  |  |  |  |
| pre-sleep | 70.18±3.59 | 69.85±4.16 | 0.09 | 0.93 |
| post-sleep | 64.83±5.38 | 61.92±4.60 | 0.52 | 0.61 |
| post- relative to pre-sleep | 90.52±4.82 | 87.19±3.69 | 0.55 | 0.59 |

**Supplementary Figure 1.**
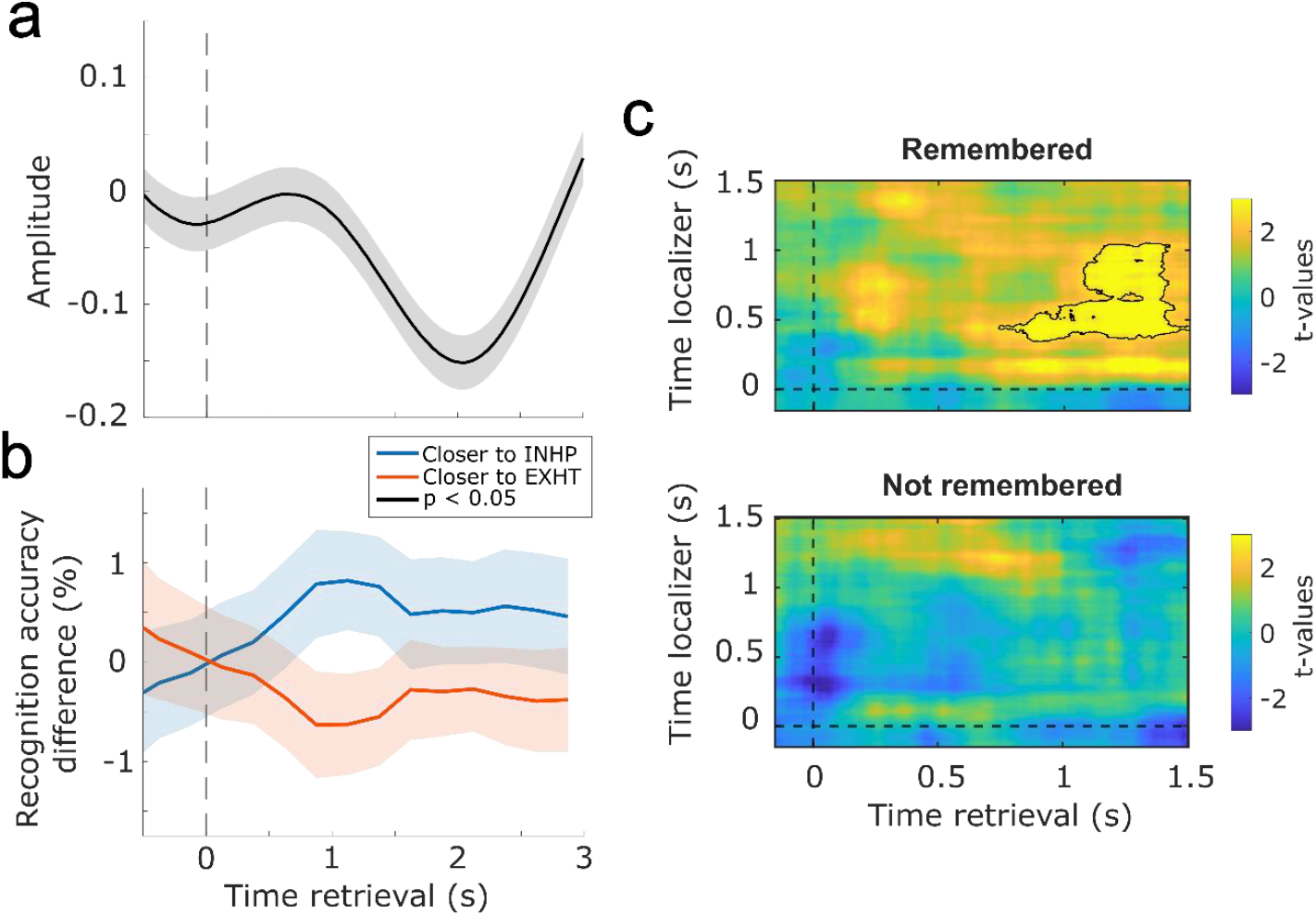
(a) Grand everage (±SEM) of the respiratory signal locked to the cue onset (t = 0 s, dashed line). The respiratory signal was recorded with a thermistor. A positive slope indicates inhalation, whereas a negative slope reflects exhalation. (b) Respiratory phase did not have any effect on recognition memory. Blue / orange lines represent the difference in accuracy with respect to the participant’s overall recognition performance for the trials that were closer to the inhalation peak / exhalation trough in each time bin (±SEM). (c) Learning-related brain patterns (objects vs. scenes) were decodable during retrieval, only for remembered trials. Contour lines indicate the extent of the significant cluster (*p* = 0.048, d = 0.7021, one-sided; corrected across times); color range (blue to yellow) represents t-values of (not) remembered against null distributions of decoding performances.

**Supplementary Figure 2.**
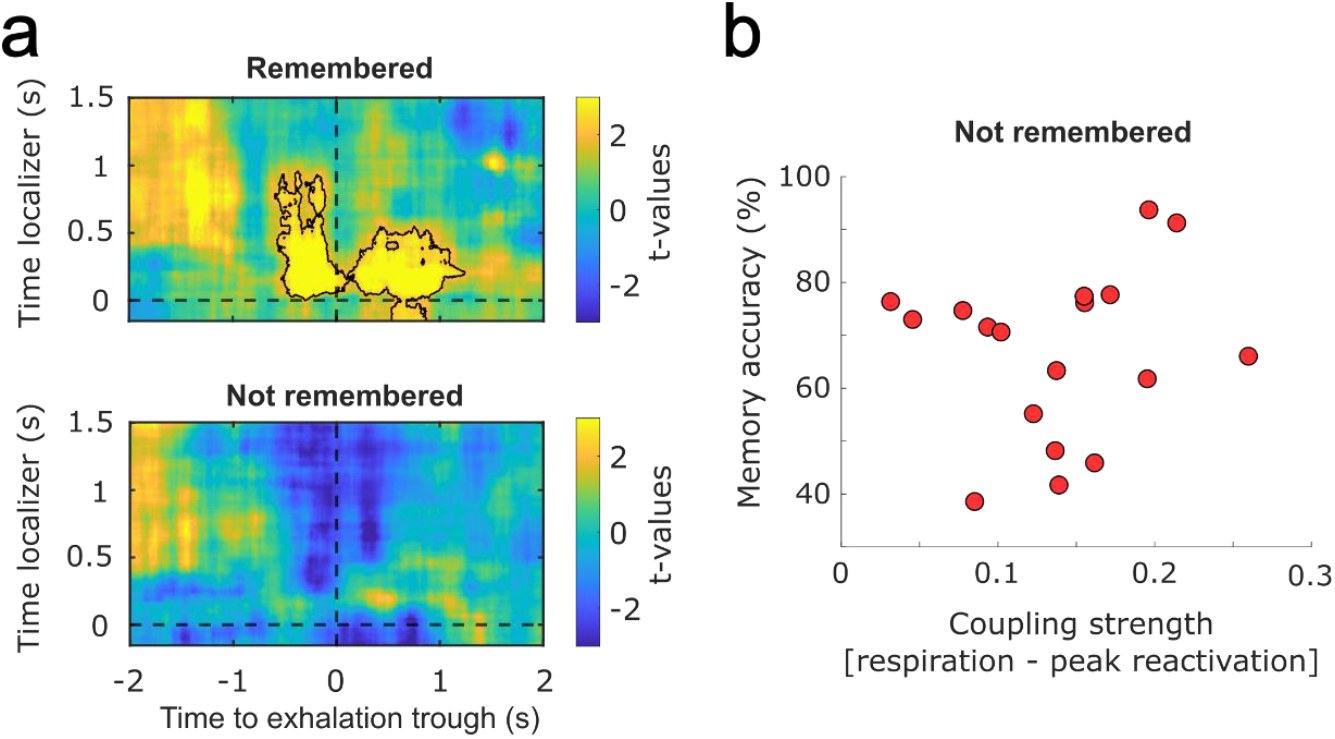
(a) Learning-related brain patterns (objects vs. scenes) were decodable during retrieval based on exhalation trough-centered EEG data, only for remembered trials. Contour lines indicate the extent of the significant cluster (p = 0.041, d = 1.0675, one-sided, corrected across times); color range (blue to yellow) represents t-values of (not) remembered against null distributions of decoding performances. (b) Coupling strength (measured as vector length) between respiration and not remembered trials’ maximum decision values from the LDA model (Spearman’s Rank Correlation, r = 0.2816, *p* = 0.2577).

